# A Suspension Trapping–Based Sample Preparation Workflow for Sensitive Plant Phosphoproteomics

**DOI:** 10.1101/2023.02.23.529696

**Authors:** Chin-Wen Chen, Chia-Feng Tsai, Miao-Hsia Lin, Shu-Yu Lin, Chuan-Chih Hsu

**Affiliations:** Institution of Plant and Microbial Biology, Academia Sinica, Taipei 115201, Taiwan; Biological Sciences Division, Pacific Northwest National Laboratory, Richland, WA 99354, United States; Department of Microbiology, College of Medicine, National Taiwan University, Taipei 100233, Taiwan; Academia Sinica Common Mass Spectrometry Facilities for Proteomics and Protein Modification Analysis, Academia Sinica, Taipei 115201, Taiwan

## Abstract

Plant phosphoproteomics provides a global view of phosphorylation-mediated signaling in plants; however, it demands high-throughput methods with sensitive detection and accurate quantification. Although protein precipitation is indispensable for removing contaminants and improving sample purity, it limits the sensitivity and throughput of plant phosphoproteomic analysis. The multiple handling steps involved in protein precipitation lead to sample loss and process variability. Herein, we developed an approach based on suspension trapping (S-Trap), termed tandem S-Trap-IMAC (immobilized metal ion affinity chromatography), by integrating an S-Trap micro column with an Fe-IMAC tip. Compared with a precipitation-based workflow, the tandem S-Trap-IMAC method deepened the coverage of the Arabidopsis (*Arabidopsis thaliana*) phosphoproteome by more than 30%, with improved quantification accuracy and short sample processing time. We applied the tandem S-Trap-IMAC method for studying abscisic acid (ABA) signaling in Arabidopsis seedlings. We thus identified 24,055 phosphopeptides and quantified several key phosphorylation sites on core ABA signaling components across four time points. Our results show that the optimized workflow aids high-throughput phosphoproteome profiling of low-input plant samples.

## INTRODUCTION

Protein phosphorylation is an important posttranslational modification for regulating plant growth and responses to environmental stress^1, 2^. For instance, upon perceiving stress signals, plants generate abscisic acid (ABA); this triggers ABA-mediated signaling pathways to activate kinases, which phosphorylate their downstream substrates, initiating defense responses such as stomatal closure and growth inhibition^3-5^. Although several core components of ABA signaling have been characterized^6, 7^, the whole signaling network is still not fully understood. This can be attributed to the complexity and dynamics of ABA-dependent phosphorylation events in plants. Therefore, systematic analysis of the entire phosphoproteome can reveal the underlying signaling networks in ABA responses^8, 9^.

Mass spectrometry (MS)–based phosphoproteomics has transformed the way phosphorylation signaling is studied in plants, enabling the comprehensive cataloging of phosphorylation events and allowing the characterization of previously unknown networks^10, 11^. However, several factors limit the efficiency of plant phosphoproteomic analysis. First, the disruption of plant cell walls requires harsh conditions and lysis buffer. Strong detergents, for instance, sodium dodecyl sulfate (SDS), have been widely employed in the lysis step to maximize the efficiency of cell wall disruption and protein extraction^12^. However, SDS suppresses proteolytic digestion and is notoriously incompatible with liquid chromatography (LC)–MS/MS analysis, reducing the detection of phosphopeptide signals^13^. Second, plant cells generate large amounts of compounds, such as photosynthetic pigments, phenolic compounds, carbohydrates, and primary and secondary metabolites, which hinders the detection of low-abundance phosphoproteins^14,15^. To address these issues, protein cleanup methods have been incorporated in sample preparation workflows to remove unwanted detergents and contaminants before proteolytic digestion^16, 17^. Currently used cleanup approaches involve protein precipitation and metabolite extraction using a combination of organic solvents such as acetone, phenol, or methanol/chloroform^18-20^. Low-mass soluble contaminants are easily separated from protein pellets through organic solvent extraction; however, inefficient resolubilization of protein pellets causes substantial sample loss and experimental variation, which is especially undesirable when plant samples are rare^21^. These challenges not only limit the depth but also decrease the quantitative accuracy of plant phosphoproteomic profiling.

Spin column–based approaches have been suggested as an alternative way of removing contaminants during plant proteomic analysis. The most common method is filter-aided sample preparation (FASP), which utilizes a 10- or 30-kDa molecular weight cutoff (MWCO) membrane to deplete small interfering substances^22^. Denatured proteins are retained on the MWCO membrane for on-filter digestion. Despite the many approaches proposed to improve the sensitivity and throughput of the FASP method^23-25^, it remains labor-intensive and leads to inefficient removal of SDS^26^. An innovative technology termed suspension trapping (S-Trap) has been developed recently for bottom-up proteomics^27^. The concept of S-Trap is to capture proteins fully denatured by SDS in a filter packed with a stack of silica fibers. Small contaminating substances can be efficiently washed off the S-Trap filter through centrifugation within minutes, significantly shortening and simplifying the sample preparation processes^28^. Notably, the enzymatic digestion step of the S-Trap procedure is completed within 1 hour, demonstrating an enhanced protease activity within the pores of the trap^29^. The S-Trap method takes advantage of the filter-based approach while addressing the problems of laborious sample preparation steps. The applicability of the S-Trap method has been shown in processing mammalian cells^30^, formalin-fixed paraffin-embedded (FFPE) tissues^31^, and urine samples^32^, achieving good proteome coverage and reproducibility. However, whether this approach can be implemented in the sample preparation workflow for plant phosphoproteomics has not yet been studied.

We established an S-Trap-based plant phosphoproteomics workflow, in which phosphopeptides are enriched using a tandem tip–based S-Trap-IMAC (immobilized metal ion affinity chromatography) platform. We carefully evaluated the influence of protein extraction buffer and S-Trap peptide elution buffer on the efficiency of phosphopeptide enrichment. We benchmarked the performance of the S-Trap-based workflow against the protein precipitation workflow in terms of phosphopeptide identification and quantification using low and high inputs of Arabidopsis (*Arabidopsis thaliana*) proteins. We further assessed the usefulness of the platform for studying the signal transduction underlying ABA treatment in Arabidopsis at four time points. The phosphoproteomics results revealed many key phosphorylation sites on the core regulators of ABA signaling across all time points, shedding light on the time-course changes of phosphorylation-mediated ABA signaling and demonstrating the feasibility of this tandem S-Trap-IMAC platform for studying plant phosphoproteomes.

## EXPERIMENTAL SECTION

### Reagents and Chemicals

Urea was purchased from Bio-Rad (Hercules, CA). Tris(2-carboxyethyl)phosphine hydrochloride (TCEP), 2-chloroacetamide (CAA), triethylammonium bicarbonate (TEAB), iron chloride, trifluoroacetic acid (TFA), ethylenediaminetetraacetic acid (EDTA), and SDS were purchased from Sigma-Aldrich (St. Louis, MO). Phosphoric acid (PA) was purchased from Honeywell Fluka (Charlotte, NC). Methanol (MeOH) and formic acid (FA) were purchased from Merck (Darmstadt, Germany). Acetic acid (AA), acetonitrile (ACN), and ammonia phosphate (NH_4_H_2_PO_4_) were purchased from J.T.Baker (Radnor, PA). Ni-NTA silica beads were purchased from Qiagen (Hilden, Germany). Empore SDB-XC solid-phase extraction disks were purchased from 3M (St. Paul, MN). S-Trap micro columns were purchased from ProtiFi (Huntington, NY). MS-grade Lys-C (lysyl endopeptidase) was purchased from FUJIFILM Wako (Osaka, Japan). Sequencing-grade modified trypsin was purchased from Promega (Madison, WI). Water was obtained from a Millipore Milli-Q system (Bedford, MA).

### Plant Culture and ABA Treatment

Sterilized Arabidopsis Columbia-0 seeds were germinated on 1/2 Murashige and Skoog (MS) medium at 4°C for 2 days. For workflow comparison, seedlings were then grown vertically in plates for 14 days and then harvested. For ABA treatment, seedlings were grown vertically in plates for 7 days and then transferred to 1/2 MS medium in a conical flask for 3 days. Seedlings were treated with or without 50 μM ABA for the times indicated.

### Protein Extraction

Arabidopsis seedlings were ground to fine powder in liquid nitrogen in a mortar. Powders were lysed in (1) 5% (w/v) SDS in 50 mM TEAB or (2) 8 M urea in 50 mM TEAB. The solution was transferred into a 1.7-mL tube and sonicated 10 times for 10 s each. The tube was centrifuged at 16,000 *g* at 4°C for 20 min, and the supernatant was collected. The protein amount was determined by bicinchoninic acid assay (Thermo Fisher Scientific, Waltham, MA).

### S-Trap and Protein Digestion

Protein digestion in the S-Trap micro column was performed according to the manufacturer’s protocol with some modifications. Briefly, 10 or 100 μg protein in lysis buffer was reduced and alkylated using 10 mM TCEP and 40 mM CAA at 45°C for 15 min. A final concentration of 5.5% (v/v) PA followed by six-fold volume of binding buffer (90%, v/v, MeOH in 100 mM TEAB) was next added to the protein solution. After gentle vortexing, the solution was loaded into an S-Trap micro column. The solution was removed by spinning the column at 4,000 *g* for 1 min. The column was washed with 150 μL binding buffer three times. Finally, 20 μL of digestion solution (1 unit Lys-C and 1 μg trypsin in 50 mM TEAB) was added to the column and incubated at 47°C for 2 h. Each digested peptide was eluted using 40 μL of three buffers consecutively: (1) 50 mM TEAB, (2) 0.2% (v/v) FA in H_2_O, and (3) 50% (v/v) ACN (standard elution). Elution solutions were collected in a tube and dried under vacuum. Peptides were desalted using an Oasis C18 cartridge (Waters). For the direct elution protocol, digests were eluted from the S-Trap micro column using 120 μL of 1% (v/v) TFA in 80% (v/v) ACN and directly loaded into a Fe-IMAC tip.

### Protein Precipitation

Protein precipitation was performed as described previously with some modifications^18^. Four volumes of MeOH followed by one volume of chloroform were added to 10 or 100 μg of lysate and mixed. Three volumes of water were then added to the tube and mixed. The solution was centrifuged at 16,000 *g* for 3 min. The upper aqueous layer was removed, the protein pellet was washed with four volumes of MeOH, and the supernatant was discarded. The precipitated protein pellet was washed again with four volumes of MeOH and then air-dried.

### Phosphopeptide Enrichment

Phosphopeptides were enriched using a Fe-IMAC tip as described previously with some modifications^33, 34^. Briefly, an IMAC tip was made in-house by plugging a 20-μm polypropylene frit disk into the tip end and packed with Ni-NTA silica resin. The packed IMAC tip was inserted into a 2-mL Eppendorf tube. Ni^2+^ ions were first removed by adding 100 mM EDTA (200 *g*, 1 min). The tip was then activated with 100 mM FeCl_3_ (200 *g*, 1 min) and equilibrated with 1% (v/v) acetic acid (200 *g*, 1 min) at pH 3.0 before sample loading. Tryptic peptides were dissolved in 1% (v/v) TFA and 80% (v/v) ACN and loaded onto the IMAC tip (200 *g*, 1 min). Next, washing steps were performed using 1% (v/v) TFA, 80% (v/v) ACN (200 *g*, 1 min), and 1% (v/v) acetic acid (pH 3.0) (200 *g*, 1 min). The IMAC tip was then inserted into an activated desalting SDB-XC StageTip. Bound phosphopeptides were eluted onto the activated desalting SDB-XC StageTip using 200 mM NH_4_H_2_PO_4_ (200 *g*, 3 min), directly eluted into sample vials for LC-MS/MS, and then dried under vacuum.

### LC-MS/MS Analysis

Dried peptides were suspended in 10 μL 0.1% (v/v) FA with 3% (v/v) ACN, and 4 μL of the sample was injected into an UltiMate 3000 UHPLC system coupled with an Orbitrap Fusion Lumos Tribrid mass spectrometer (Thermo Fisher Scientific). Buffer A was 0.1% (v/v) FA, and buffer B was 0.1% (v/v) FA in 100% ACN. The LC gradient over 80 min was established as follows: 2 min from 1.6% to 5% buffer B, 60 min from 5% to 28% buffer B, 6 min from 28% to 50% buffer B, 1 min from 50% to 95% buffer B, 5 min of 95% buffer B, 1 min from 95% to 1.6% buffer B, and 5 min of 1.6% buffer B. Peptides were separated on a 25-cm Thermo Acclaim PepMap RSLC column with a column heater set at 45°C. The mass spectrometer was operated in data-dependent acquisition mode, in which a full MS scan (m/z 375–1,600; resolution: 60,000) was followed by the most intense ions being subjected to higher-energy collision-induced dissociation (HCD) fragmentation within 3 s. Data were acquired in the Orbitrap with a resolution of 15,000 [normalized collision energy (NCE): 30%; max injection time: 100 ms; isolation window: 1.4 m/z; dynamic exclusion: 20 s].

### Data Processing

MS raw files were searched against the TAIR10 database (70,774 entries with 50% decoys) using the MSFragger^35^ (version 3.4) search engine of the FragPipe platform (version 18.0). The LFQ-phospho workflow in FragPipe was selected with some modifications. Precursor mass tolerance was set to ±10 ppm, and fragment mass tolerance was set to 20 ppm. Protein digestion was set to trypsin (after KR/-) up to two missed cleavages. The fixed modification was set as carbamidomethyl (C), and variable modifications were set as oxidation (M), acetylation (protein N-term), and phosphorylation (STY) for phosphoproteomic analysis. The match-between-runs (MBR) function was enabled with 1% MBR ion false discovery rate (FDR) and 1 min MBR retention time (RT) tolerance. The FDR for peptide-spectrum match (PSM), ion, peptide, and protein level was set at 1%.

### Data Analysis

The number of unique phosphopeptides identified from each sample was calculated using Microsoft Excel in the FragPipe output combined_modified_peptide table. A localization probability of 0.75 was used as the cutoff for confidence of phosphorylation sites. The intensities of all phosphorylation sites identified were extracted from the FragPipe output STY_79.9663 table for further quantitative analysis. Perseus software^36^ (version 1.6.15.0) was used for data and statistical analysis. The intensities of localized phosphorylation sites were log2 transformed, and quantifiable phosphorylation sites were selected from the site intensity of all triplicate replicates in at least one ABA treatment group. Pearson correlation of phosphopeptide intensity between replicates was calculated by Perseus software. Missing intensities were imputed from a normal distribution (width = 0.2 and downshift = 2.0) for ANOVA test. ANOVA test was performed for selecting significantly enriched phosphorylation sites using a permutation-based FDR threshold of 0.05 and S_0_ = 0.5. Principal component analysis (PCA) of the ABA-treated samples was performed using phosphorylation sites quantified across 16 samples with a Benjamini–Hochberg FDR cutoff of 0.05. For heatmaps, the mean intensities from triplicate replicates of each phosphorylation site were z-scored. Heatmaps were generated using the color scale in Microsoft Excel.

### Data Availability

The MS raw data and FragPipe output files have been deposited with the ProteomeXchange Consortium via the jPOST^37^ repository with the data set identifier PXD039933.

## RESULTS AND DISCUSSION

### Tandem S-Trap-IMAC Phosphoproteomics Workflow

In conventional plant phosphoproteomics, protein precipitation and peptide desalting are critical steps in sample preparation to eliminate the interference of low-mass contaminants in LC-MS/MS detection. However, laborious steps and considerable sample loss during protein precipitation limit the throughput and depth of plant phosphoproteome analysis. To address this issue, we developed a streamlined tandem S-Trap-IMAC workflow for plant phosphoproteomics to avoid contaminant interference without the need for protein precipitation and to enrich phosphopeptides by direct loading of digested peptides from the S-Trap into the Fe-IMAC tip (Figure 1). This workflow is based on the concept that small interfering contaminants can pass through the silica fibers in the S-Trap column while proteins are captured, allowing S-Trap proteolytic digestion. We then used an optimized buffer for simultaneous elution of S-Trap peptides and binding of Fe-IMAC phosphopeptides. Phosphorylated peptides were isolated by centrifuging the tandem S-Trap-IMAC tip to minimize sample loss caused by solution transfer and tube collection.

**Figure 1.**
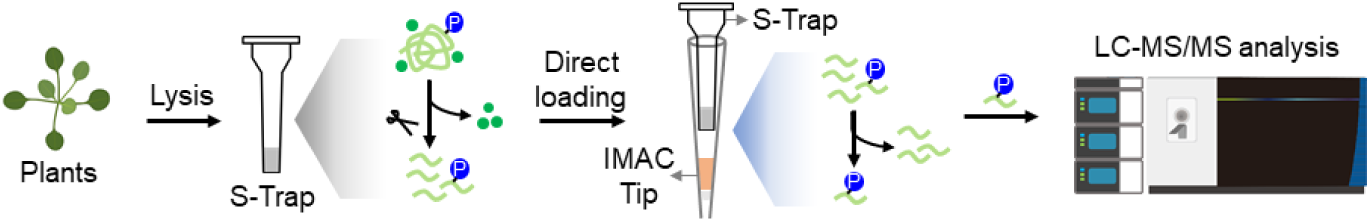
Experimental design of the tandem S-Trap-IMAC workflow for plant phosphoproteomics. Interfering contaminants in the plant lysate are removed by washing the S-Trap, and proteins are digested within the S-Trap. Phosphopeptides are enriched using an Fe-IMAC tip via a direct loading strategy and analyzed using a Fusion Lumos mass spectrometer in data-dependent acquisition mode.

### Efficiency of Protein Extraction Buffers for Plant Tissues

Recently, we reported a sample preparation workflow using guanidine hydrochloride (GdnHCl) as a lysis reagent for plant tissues^16^. However, GdnHCl is not compatible with the S-Trap workflow because positively charged GdnHCl precipitates with negatively charged SDS. To our knowledge, there is no literature reporting the protein yield of different lysis strategies for plant tissues using the S-Trap method. To determine the efficiency of lysis buffers for our S-Trap-based workflow, we tested two protein extraction protocols: (1) SDS and (2) Urea-SDS (Figure S1A). Proteins were first extracted using 5% (w/v) SDS (SDS protocol) or 8 M urea (Urea-SDS protocol). Next, 100 μg of proteins extracted using 8 M urea was further denatured by adding the same volume of 10% (w/v) SDS (Urea-SDS protocol), and 100 μg of denatured proteins from the two protocols was digested in S-Trap columns. Digested phosphopeptides were eluted into tubes and dried under vacuum, following the standard S-Trap protocol. Dried phosphopeptides were suspended in IMAC loading buffer, enriched using a Fe-IMAC tip, and analyzed by nano-LC-MS/MS.

Surprisingly, the number of phosphopeptides identified was significantly increased (*p* value < 0.05) from 4,728 and 4,902 using the SDS protocol to 5,391 and 6,215 using the Urea-SDS protocol, corresponding to a 14% and 27% improvement using MS/MS or MBR, respectively (Figure 2A). To determine the reason for performance improvement, we compared the overlap of phosphopeptides identified using the two protocols (Figure S1B). The 5,956 phosphopeptides identified using both protocols displayed the same intensity distribution (Figure 2B), suggesting that the two protocols have similar extraction efficiency for highly abundant phosphoproteins. By contrast, the median phosphopeptide intensity of the 3,175 phosphopeptides identified using only the Urea-SDS protocol was lower than that of the 2,196 phosphopeptides identified using only the SDS protocol (Figure 2C). The lower median phosphopeptide intensity from the Urea-SDS protocol indicates that the protocol identified more low-abundance phosphopeptides, suggesting that it may extract more phosphoproteins present in low quantities than the SDS protocol. We therefore chose the Urea-SDS protocol for the protein extraction steps in the S-Trap workflow.

**Figure 2.**
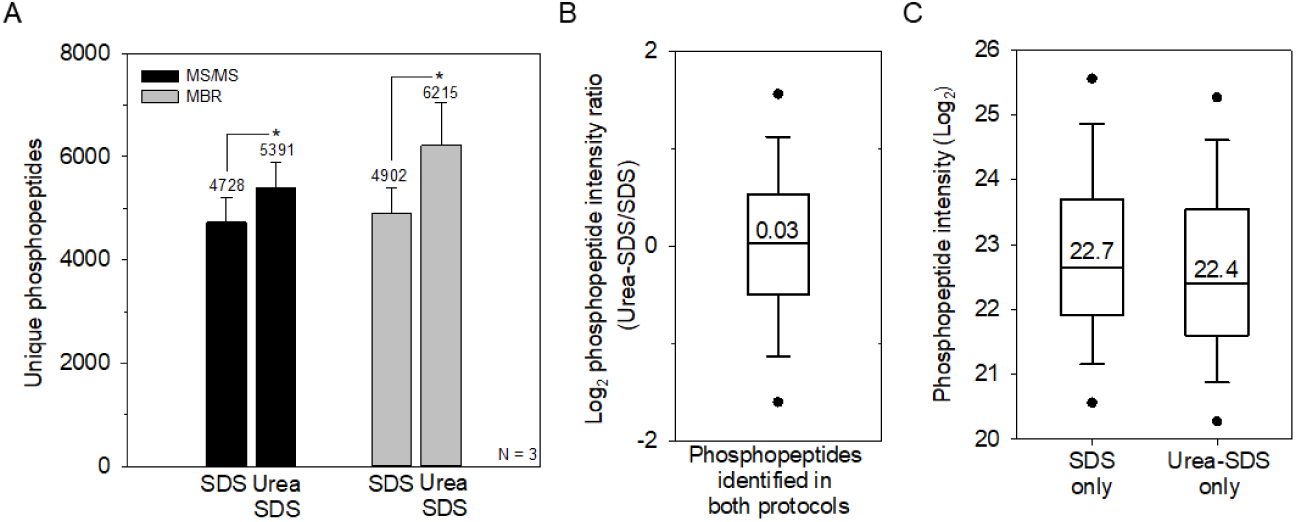
Extraction efficiency for plant phosphoproteins using the S-Trap workflow. (A) Number of unique phosphopeptides using the SDS and Urea-SDS protocols. Error bars, standard deviation (SD) (n = 3). Two-Sample t-test, \**p* < 0.05. (B) Log2-transformed intensity ratio of the phosphopeptides identified in both protocols. Boxes mark the first, median, and third quantiles, and whiskers mark the minimum/maximum value within 1.5 interquartile range. Outliers are not shown. (C) Distribution of the log2-transformed intensity of the phosphopeptides identified using either protocol. Boxes mark the first, median, and third quantiles, and whiskers mark the minimum/maximum value within 1.5 interquartile range. Outliers are not displayed.

### Tandem S-Trap-IMAC Strategy for Plant Phosphoproteomics

Since the elution buffer for releasing digested peptides from the S-Trap is not compatible with the Fe-IMAC, tube collection and buffer exchange steps are required before phosphopeptide enrichment. To improve the reproducibility and throughput of phosphopeptide enrichment, we designed a tandem S-Trap-IMAC workflow, integrating an S-Trap micro column and Fe-IMAC tip for direct loading of digested peptides from the S-Trap into the Fe-IMAC tip (direct loading protocol). We therefore evaluated the use of a compatible buffer for both S-Trap peptide elution and IMAC loading. We hypothesized that high-concentration (80%) ACN with 1% TFA would efficiently elute peptides from the S-Trap without affecting the enrichment specificity of the Fe-IMAC. To test this hypothesis, we compared the performance of the standard S-Trap elution protocol and the direct loading protocol (Figure S2A). Compared with the standard elution protocol, the direct loading one slightly improved the number of phosphopeptides identified (from 6,215 to 6,813 in MBR), which suggests that both protocols offer similar peptide elution efficiency (Figure 3A).

**Figure 3.**
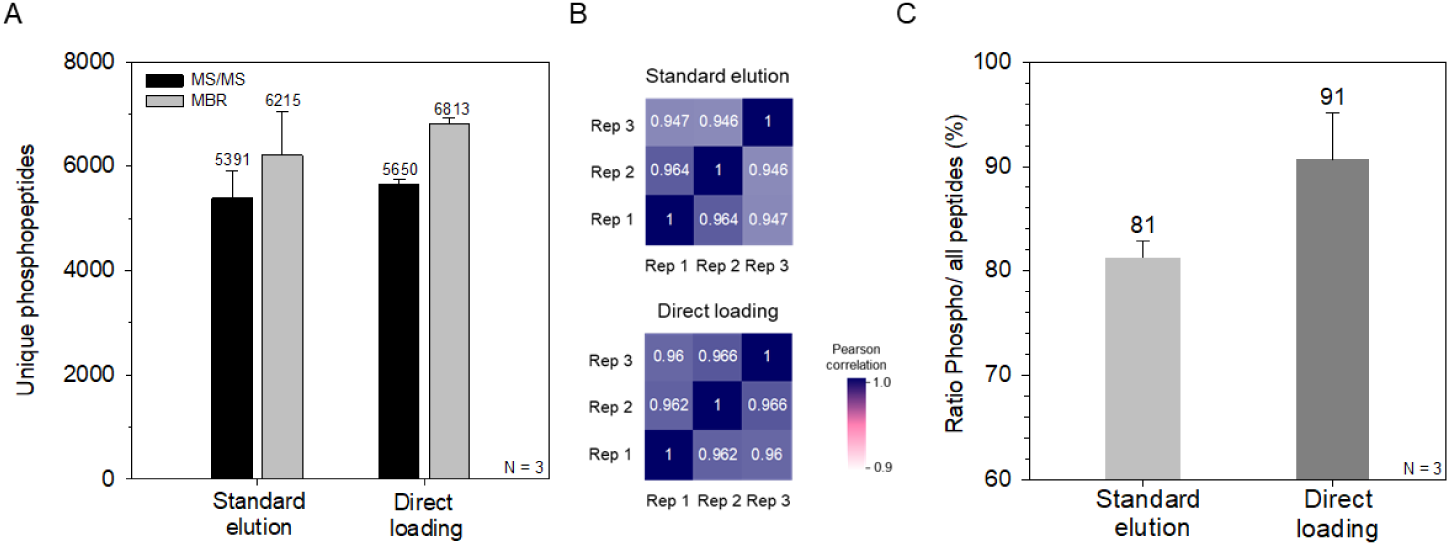
Performance of standard elution and direct loading protocols. (A) Number of phosphopeptides identified via MS/MS and MBR methods. Error bars, SD (n = 3). (B) Pearson correlation of phosphopeptide intensity between triplicate replicates. (C) Enrichment selectivity of the Fe-IMAC. Error bars, SD (n = 3).

To assess the robustness of the two protocols, we compared the Pearson correlations of log2-transformed phosphopeptide intensities for both of them. For the standard elution protocol, correlation coefficients between replicates ranged from 0.946 to 0.964. By contrast, high coefficient values and a narrow coefficient range (0.96 to 0.966) were observed for the direct loading protocol (Figure 3B). In addition, the direct loading protocol generated a higher percentage of quantifiable phosphopeptides (<30% missing values in three replicates) than the standard elution protocol (Figure S2B). From these results, we determined that the direct loading protocol provides better reproducibility relative to the standard S-Trap elution protocol. We also evaluated whether the direct loading strategy affects Fe-IMAC enrichment specificity. More than 90% of the peptides identified using the direct loading protocol were phosphorylated, which was higher than the mean percentage (81%) obtained using the standard elution protocol (Figure 3C). This demonstrates that 80% ACN in 1% TFA is an ideal buffer for the direct loading strategy.

### Tandem S-Trap-IMAC Strategy Outperforms Protein Precipitation–Based Workflow

We next evaluated whether the tandem S-Trap-IMAC workflow outperforms conventional workflows based on protein precipitation in terms of sensitivity, reproducibility, and throughput. For this purpose, we performed low-input (10 μg protein) and high-input (100 μg protein) Arabidopsis phosphoproteomics using tandem S-Trap-IMAC (S-Trap protocol) and protein precipitation (precipitation protocol) sample preparation workflows, respectively (Figure S3A). The S-Trap protocol allowed the identification of 6,813 and 601 phosphopeptides from 100 and 10 μg protein input, respectively, using the MBR function across three replicates (Figures 4A and 4B). This was an improvement of more than 30% compared with the precipitation protocol with both low and high protein inputs, with a *p* value less than 0.05, suggesting that the S-Trap protocol reduces sample loss by skipping the protein precipitation steps. To examine whether the improvement in the number of phosphopeptides identified using the S-Trap protocol was related to the molecular weight (MW) and isoelectric point (pI) of the phosphoproteins, we compared the distributions of phosphoprotein MW and pI in each protocol. Lack of distinct patterns between the two protocols suggests that the improvement might have resulted from reduced loss of phosphoproteins on the tube surface rather than the protein precipitation and resolubilization steps (Figures 4C and 4D).

**Figure 4.**
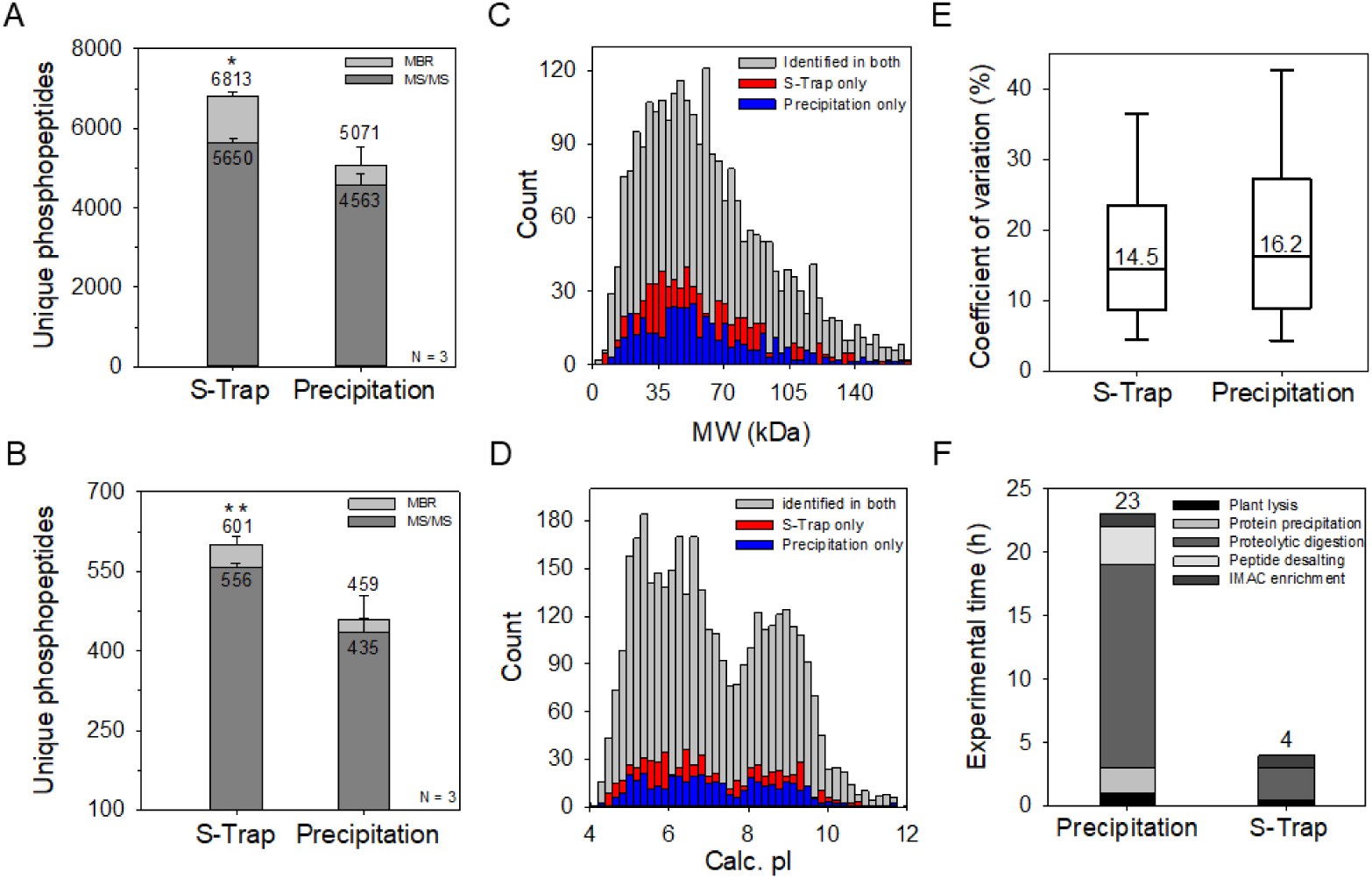
Comparison of plant phosphoproteomics workflows based on S-Trap-IMAC and protein precipitation. Number of phosphopeptides identified using (A) high-input and (B) low-input protocols. Error bars, SD (n = 3). Two-Sample t-test, \**p* < 0.05, ** *p* < 0.01. Distribution of (C) molecular weight and (D) isoelectric point of phosphoproteins identified. (E) Distribution of coefficient of variation (%) of triplicate replicates. Boxes mark the first, median, and third quantiles, and whiskers mark the minimum/maximum value within 1.5 interquartile range. Outliers are not displayed. (F) Total experimental time of the two protocols.

In addition to the number of phosphopeptides identified, we also calculated the coefficients of variation (CVs) for the two protocols (Figure 4E). The median CV was 14.5% in the S-Trap protocol, whereas the corresponding CV was 16.2% in the precipitation protocol. The multiple buffer exchange steps and protein precipitation and resolubilization steps in the protein precipitation workflow may be the main cause of the higher CV in the precipitation protocol. The total sample processing time for the S-Trap protocol was 4 h, compared with approximately 1 day for the precipitation protocol, which is attributed to the simplified sample preparation steps and shortened proteolytic time (2 h) in the S-Trap system (Figure 4F). Notably, the percentage of miscleaved phosphopeptides (34%) in the S-Trap protocol was lower than that in the precipitation protocol (40%), demonstrating that the digestion efficiency of the S-Trap protocol is superior to that of the precipitation protocol even when overnight digestion is performed in the precipitation protocol (Figure S3B). Overall, we demonstrated that the tandem S-Trap-IMAC strategy outperforms the conventional precipitation-based workflow in terms of sensitivity, reproducibility, and throughput.

### Application of S-Trap-IMAC for Studying ABA-Mediated Signaling in Arabidopsis

To demonstrate the scalability of the tandem S-Trap-IMAC method for plant phosphoproteomic analysis, we applied it to phosphoproteome profiling of ABA-treated Arabidopsis. We treated Arabidopsis with 50 μM ABA for 0, 10, 30, or 60 min and prepared three biological replicates for each time point (Figure 5A). After ABA treatment, 100 μg protein input was used for S-Trap digestion, S-Trap-IMAC enrichment, and nano-LC-MS/MS analysis. Our analysis identified 24,055 unique phosphopeptides in 16 samples, corresponding to 4,147 phosphoproteins (Tables S1 and S2). Approximately 7,500 unique phosphopeptides were identified in each replicate (Figure S4). We were able to localize 9,867 phosphorylation sites (probability > 75%) on the phosphopeptides identified, with 4,626 phosphorylation sites quantifiable using a criterion of less than 30% missing data. To evaluate the sample quality and consistency for each treatment, we performed PCA based on the phosphorylation sites identified in all replicates. The four treatment groups were clearly separated, while the biological replicates at each time point were close together (Figure 5B). The PCA results indicated that ABA changes the phosphoproteomic profile of Arabidopsis in a time-dependent manner. We next performed an ANOVA test with a 5% FDR cutoff to identify phosphorylation sites significantly perturbed after ABA treatment. The test indicated that 57% (2,617) of quantifiable phosphorylation sites were differentially phosphorylated between different time points. Notably, 857 phosphorylation sites were changed after 10 min of ABA treatment, implying that ABA induces phosphorylation events early in Arabidopsis.

**Figure 5.**
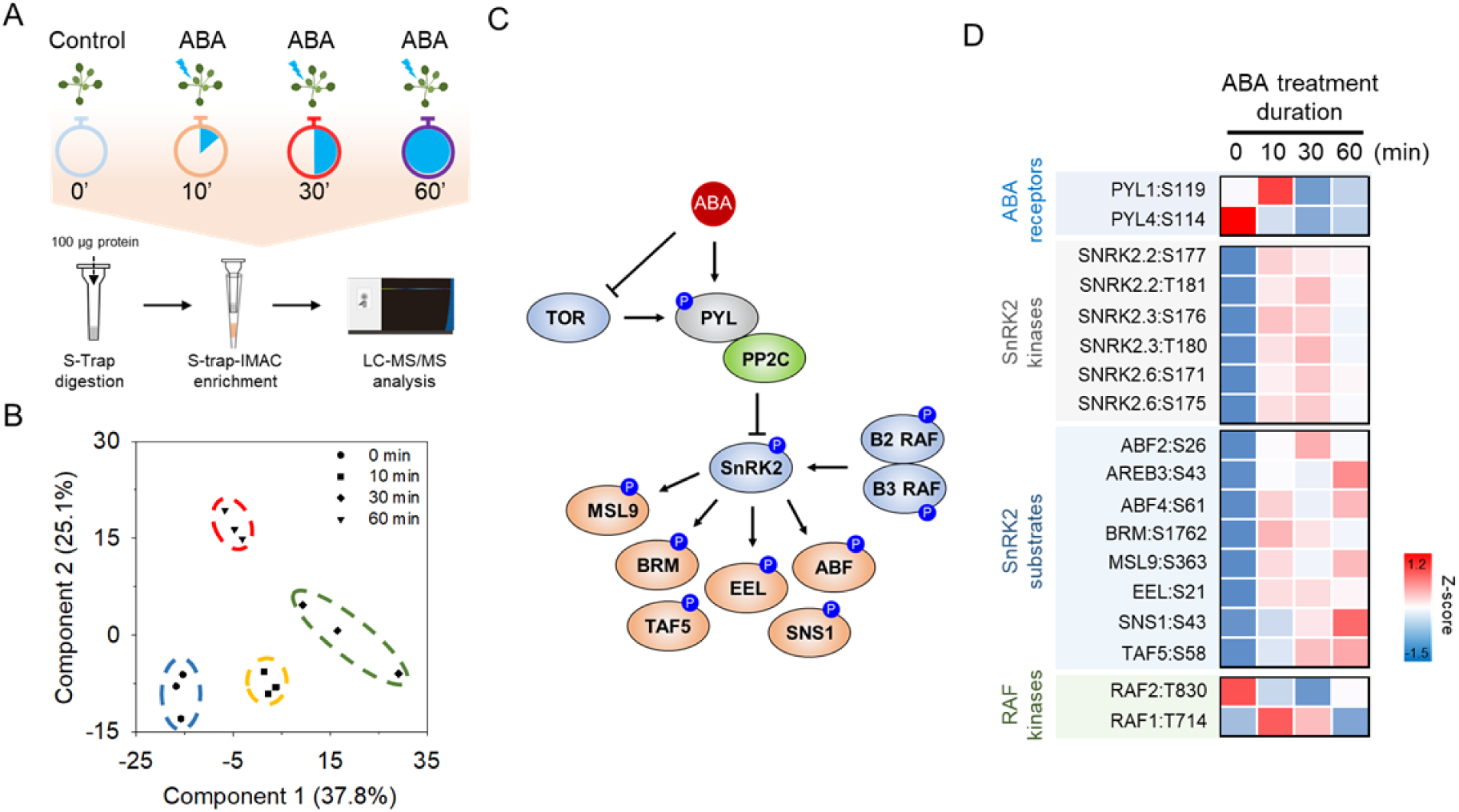
Phosphoproteomic analysis of ABA-dependent signaling in Arabidopsis using the S-Trap-IMAC workflow. (A) Schematic representation of the experimental workflow. (B) PCA of phosphorylation sites across all replicates. (C) Overview of ABA signaling components identified in Arabidopsis. (D) Heatmap showing the relative intensity of key phosphorylation sites during time-dependent ABA signaling.

Our phosphoproteomic analysis covered many key components of ABA signaling in Arabidopsis (Figure 5C). In the ABA core signaling pathway, ABA receptors Pyrabactin Resistance 1 (PYR1)/PYLs/RCARs bind to ABA and then interact with PP2C phosphatases, leading to release of SnRK2 kinases from PP2C-mediated inhibition^38-41^. Raf-like kinases (RAFs) then phosphorylate SnRK2s at the activation loop, triggering SnRK2 activation^42-44^. Activated SnRK2s further phosphorylate downstream substrates to induce plant stress responses^45, 46^. In addition, the ABA binding activity of PYR1/PYLs/RCARs is regulated by Target of Rapamycin (TOR) kinases^47^. Two TOR phosphorylation sites of PYLs decreased the phosphorylation level after 10 min of ABA treatment (Figure 5D and Table S3). In addition, we did not observe a significant increase in phosphorylation of two RAF kinases during ABA treatment. This is consistent with literature reporting that ABA does not increase RAF activity and that RAFs only require basal activity to phosphorylate and initiate the autophosphorylation of SnRK2 kinases^44^. Phosphorylation of the activation loops of SnRK2s was detected at 10 min and peaked at 30 min of ABA treatment, suggesting the quick activation of SnRK2s in ABA signaling. Finally, most direct substrates of SnRK2s, such as AREB3^45^, ABF4^45^, SNS1^8^, and MSL9^9^, showed the highest phosphorylation levels after 1 h of ABA treatment, suggesting that these are indeed the downstream layer of ABA signaling after SnRK2 activation. The tandem S-Trap-IMAC approach was able to monitor phosphorylation changes of core phosphoproteins involved in ABA signaling during a time-course experiment, demonstrating that this strategy may be applicable for studying other events mediated by phosphorylation in plants.

## CONCLUSION

We successfully integrated an S-Trap micro column with a Fe-IMAC tip to directly enrich phosphopeptides after digestion in the S-Trap without the need for buffer exchange or tube collection. The use of 8 M urea with the SDS extraction buffer enhanced protein extraction efficiency during plant lysis, while the replacement of multiple elution buffers in the S-Trap protocol with a single IMAC-compatible elution buffer did not affect the selectivity of Fe-IMAC. This direct loading strategy significantly improved the reproducibility of S-Trap-based plant phosphoproteomics.

Our findings reveal that the tandem S-Trap-IMAC strategy outperformed conventional workflows based on protein precipitation in terms of phosphopeptide identification, quantification accuracy, and throughput. This improvement is likely due to simplifying the sample preparation steps before phosphopeptide enrichment, thereby reducing sample loss, in the tandem S-Trap-IMAC workflow. We systematically analyzed the effects of the important phytohormone ABA on the Arabidopsis phosphoproteome using the tandem S-Trap-IMAC approach, and the results demonstrated great coverage of the core components of ABA signaling. This suggests that the S-Trap-IMAC approach could be applied to large-scale signaling studies of mass-limited samples in a high-throughput manner, which remains unattainable using current plant phosphoproteomics workflows.

## ASSOCIATED CONTENT

### Supporting Information

Figure S1: Comparison of the overlap of identified phosphopeptides from the SDS protocol and the Urea-SDS protocol; Figure S2: Comparison of the ratio of quantifiable phosphopeptides from the standard elution protocol and the direct loading protocol; Figure S3: Comparison of the ratio of missed cleavage from the S-Trap protocol and the precipitation protocol; Figure S4: The identified phosphopeptides of each replicate; Table S1: The total number of identified phosphoproteins, phosphopeptides, and phosphorylation sites in the ABA-dependent phosphoproteomics experiment; Table S2: Identified phosphopeptides and their intensities and localization probabilities in the ABA-dependent phosphoproteomics experiment (XLSX); Table S3: The z-scored intensities of selected ABA signaling components (XLSX).

The Supporting Information is available free of charge on the ACS Publications website.

## AUTHOR INFORMATION

### Author Contributions

C.C.H. designed the experiments. C.W.C. and S.Y.L. performed experiments. C.W.C., C.F.T. and C.C.H analyzed data. M.H.L. and C.C.H. wrote the manuscript. All authors have given approval to the final version of the manuscript.

### Note

The authors declare no financial interest.

## ACKNOWLEDGMENTS

This research was supported by a grant (110-2311-B-001-043-MY2) from MOST and Academia Sinica Core Facility and Innovative Instrument Project (AS-CFII-111-209) from Academia Sinica. Mass spectrometry data were acquired at the Medicinal Chemistry and Analytical Core Facilities in the Biomedical Translation Research Center located at National Biotechnology Research Park, supported by Academia Sinica Core Facility and Innovative Instrument (AS-NBRPCF-111-201). The research was performed in the Proteomics Core Lab, Institute of Plant and Microbial Biology, Academia Sinica.

## Table of Contents

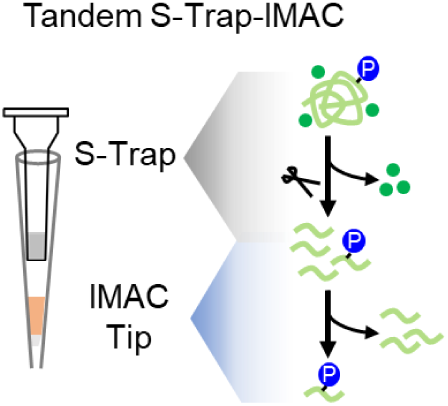

## REFERENCES

1. Park, C. J.; Caddell, D. F.; Ronald, P. C., Protein phosphorylation in plant immunity: insights into the regulation of pattern recognition receptor-mediated signaling. Front Plant Sci 2012, 3, 177.

2. Damaris, R. N.; Yang, P., Protein Phosphorylation Response to Abiotic Stress in Plants. Methods Mol Biol 2021, 2358, 17–43.

3. Umezawa, T.; Takahashi, F.; Shinozaki, K., Phosphorylation networks in the abscisic Acid signaling pathway. Enzymes 2014, 35, 27–56.

4. Zhu, J. K., Abiotic Stress Signaling and Responses in Plants. Cell 2016, 167 (2), 313–324.

5. Chen, K.; Li, G. J.; Bressan, R. A.; Song, C. P.; Zhu, J. K.; Zhao, Y., Abscisic acid dynamics, signaling, and functions in plants. J Integr Plant Biol 2020, 62 (1), 25–54.

6. Cutler, S. R.; Rodriguez, P. L.; Finkelstein, R. R.; Abrams, S. R., Abscisic acid: emergence of a core signaling network. Annu Rev Plant Biol 2010, 61, 651–79.

7. Raghavendra, A. S.; Gonugunta, V. K.; Christmann, A.; Grill, E., ABA perception and signalling. Trends Plant Sci 2010, 15 (7), 395–401.

8. Umezawa, T.; Sugiyama, N.; Takahashi, F.; Anderson, J. C.; Ishihama, Y.; Peck, S. C.; Shinozaki, K., Genetics and phosphoproteomics reveal a protein phosphorylation network in the abscisic acid signaling pathway in Arabidopsis thaliana. Sci Signal 2013, 6 (270), rs8.

9. Wang, P.; Xue, L.; Batelli, G.; Lee, S.; Hou, Y. J.; Van Oosten, M. J.; Zhang, H.; Tao, W. A.; Zhu, J. K., Quantitative phosphoproteomics identifies SnRK2 protein kinase substrates and reveals the effectors of abscisic acid action. Proc Natl Acad Sci U S A 2013, 110 (27), 11205–10.

10. Kersten, B.; Agrawal, G. K.; Durek, P.; Neigenfind, J.; Schulze, W.; Walther, D.; Rakwal, R., Plant phosphoproteomics: an update. Proteomics 2009, 9 (4), 964–88.

11. Silva-Sanchez, C.; Li, H.; Chen, S., Recent advances and challenges in plant phosphoproteomics. Proteomics 2015, 15 (5-6), 1127–41.

12. Chen, E. I.; Cociorva, D.; Norris, J. L.; Yates, J. R., 3rd, Optimization of mass spectrometry-compatible surfactants for shotgun proteomics. J Proteome Res 2007, 6 (7), 2529–38.

13. Min, L.; Choe, L. H.; Lee, K. H., Improved protease digestion conditions for membrane protein detection. Electrophoresis 2015, 36 (15), 1690–8.

14. Isaacson, T.; Damasceno, C. M.; Saravanan, R. S.; He, Y.; Catala, C.; Saladie, M.; Rose, J. K., Sample extraction techniques for enhanced proteomic analysis of plant tissues. Nat Protoc 2006, 1 (2), 769–74.

15. Wu, X.; Gong, F.; Wang, W., Protein extraction from plant tissues for 2DE and its application in proteomic analysis. Proteomics 2014, 14 (6), 645–58.

16. Hsu, C. C.; Zhu, Y.; Arrington, J. V.; Paez, J. S.; Wang, P.; Zhu, P.; Chen, I. H.; Zhu, J. K.; Tao, W. A., Universal Plant Phosphoproteomics Workflow and Its Application to Tomato Signaling in Response to Cold Stress. Mol Cell Proteomics 2018, 17 (10), 2068–2080.

17. Song, G.; Hsu, P. Y.; Walley, J. W., Assessment and Refinement of Sample Preparation Methods for Deep and Quantitative Plant Proteome Profiling. Proteomics 2018, 18 (17), e1800220.

18. Wessel, D.; Flugge, U. I., A method for the quantitative recovery of protein in dilute solution in the presence of detergents and lipids. Anal Biochem 1984, 138 (1), 141–3.

19. Damerval, C.; Devienne, D.; Zivy, M.; Thiellement, H., Technical Improvements in Two-Dimensional Electrophoresis Increase the Level of Genetic-Variation Detected in Wheat-Seedling Proteins. Electrophoresis 1986, 7 (1), 52–54.

20. Hurkman, W. J.; Tanaka, C. K., Solubilization of Plant Membrane-Proteins for Analysis by Two-Dimensional Gel-Electrophoresis. Plant Physiol 1986, 81 (3), 802–806.

21. Mergner, J.; Kuster, B., Plant Proteome Dynamics. Annu Rev Plant Biol 2022, 73, 67–92.

22. Wisniewski, J. R.; Zougman, A.; Nagaraj, N.; Mann, M., Universal sample preparation method for proteome analysis. Nat Methods 2009, 6 (5), 359–62.

23. Erde, J.; Loo, R. R.; Loo, J. A., Enhanced FASP (eFASP) to increase proteome coverage and sample recovery for quantitative proteomic experiments. J Proteome Res 2014, 13 (4), 1885–95.

24. Huber, M. L.; Sacco, R.; Parapatics, K.; Skucha, A.; Khamina, K.; Muller, A. C.; Rudashevskaya, E. L.; Bennett, K. L., abFASP-MS: affinity-based filter-aided sample preparation mass spectrometry for quantitative analysis of chemically labeled protein complexes. J Proteome Res 2014, 13 (2), 1147–55.

25. Glatter, T.; Ahrne, E.; Schmidt, A., Comparison of Different Sample Preparation Protocols Reveals Lysis Buffer-Specific Extraction Biases in Gram-Negative Bacteria and Human Cells. J Proteome Res 2015, 14 (11), 4472–85.

26. Mikulasek, K.; Konecna, H.; Potesil, D.; Holankova, R.; Havlis, J.; Zdrahal, Z., SP3 Protocol for Proteomic Plant Sample Preparation Prior LC-MS/MS. Front Plant Sci 2021, 12, 635550.

27. Zougman, A.; Selby, P. J.; Banks, R. E., Suspension trapping (STrap) sample preparation method for bottom-up proteomics analysis. Proteomics 2014, 14 (9), 1006–0.

28. HaileMariam, M.; Eguez, R. V.; Singh, H.; Bekele, S.; Ameni, G.; Pieper, R.; Yu, Y., S-Trap, an Ultrafast Sample-Preparation Approach for Shotgun Proteomics. J Proteome Res 2018, 17 (9), 2917–2924.

29. Hayoun, K.; Gouveia, D.; Grenga, L.; Pible, O.; Armengaud, J.; Alpha-Bazin, B., Evaluation of Sample Preparation Methods for Fast Proteotyping of Microorganisms by Tandem Mass Spectrometry. Front Microbiol 2019, 10, 1985.

30. Bettinger, J. Q.; Welle, K. A.; Hryhorenko, J. R.; Ghaemmaghami, S., Quantitative Analysis of in Vivo Methionine Oxidation of the Human Proteome. J Proteome Res 2020, 19 (2), 624–633.

31. Marchione, D. M.; Ilieva, I.; Devins, K.; Sharpe, D.; Pappin, D. J.; Garcia, B. A.; Wilson, J. P.; Wojcik, J. B., HYPERsol: High-Quality Data from Archival FFPE Tissue for Clinical Proteomics. J Proteome Res 2020, 19 (2), 973–983.

32. Ding, H.; Fazelinia, H.; Spruce, L. A.; Weiss, D. A.; Zderic, S. A.; Seeholzer, S. H., Urine Proteomics: Evaluation of Different Sample Preparation Workflows for Quantitative, Reproducible, and Improved Depth of Analysis. J Proteome Res 2020, 19 (4), 1857–1862.

33. Tsai, C. F.; Hsu, C. C.; Hung, J. N.; Wang, Y. T.; Choong, W. K.; Zeng, M. Y.; Lin, P. Y.; Hong, R. W.; Sung, T. Y.; Chen, Y. J., Sequential phosphoproteomic enrichment through complementary metal-directed immobilized metal ion affinity chromatography. Anal Chem 2014, 86 (1), 685–93.

34. Tsai, C. F.; Wang, Y. T.; Hsu, C. C.; Kitata, R. B.; Chu, R. K.; Velickovic, M.; Zhao, R.; Williams, S. M.; Chrisler, W. B.; Jorgensen, M. L.; Moore, R. J.; Zhu, Y.; Rodland, K. D.; Smith, R. D.; Wasserfall, C. H.; Shi, T.; Liu, T., A streamlined tandem tip-based workflow for sensitive nanoscale phosphoproteomics. Commun Biol 2023, 6 (1), 70.

35. Kong, A. T.; Leprevost, F. V.; Avtonomov, D. M.; Mellacheruvu, D.; Nesvizhskii, A. I., MSFragger: ultrafast and comprehensive peptide identification in mass spectrometry-based proteomics. Nat Methods 2017, 14 (5), 513–520.

36. Tyanova, S.; Temu, T.; Sinitcyn, P.; Carlson, A.; Hein, M. Y.; Geiger, T.; Mann, M.; Cox, J., The Perseus computational platform for comprehensive analysis of (prote)omics data. Nat Methods 2016, 13 (9), 731–40.

37. Moriya, Y.; Kawano, S.; Okuda, S.; Watanabe, Y.; Matsumoto, M.; Takami, T.; Kobayashi, D.; Yamanouchi, Y.; Araki, N.; Yoshizawa, A. C.; Tabata, T.; Iwasaki, M.; Sugiyama, N.; Tanaka, S.; Goto, S.; Ishihama, Y., The jPOST environment: an integrated proteomics data repository and database. Nucleic Acids Res 2019, 47 (D1), D1218–D1224.

38. Yoshida, R.; Hobo, T.; Ichimura, K.; Mizoguchi, T.; Takahashi, F.; Aronso, J.; Ecker, J. R.; Shinozaki, K., ABA-activated SnRK2 protein kinase is required for dehydration stress signaling in Arabidopsis. Plant Cell Physiol 2002, 43 (12), 1473–83.

39. Fujii, H.; Zhu, J. K., Arabidopsis mutant deficient in 3 abscisic acid-activated protein kinases reveals critical roles in growth, reproduction, and stress. Proc Natl Acad Sci U S A 2009, 106 (20), 8380–5.

40. Ma, Y.; Szostkiewicz, I.; Korte, A.; Moes, D.; Yang, Y.; Christmann, A.; Grill, E., Regulators of PP2C phosphatase activity function as abscisic acid sensors. Science 2009, 324 (5930), 1064–8.

41. Park, S. Y.; Fung, P.; Nishimura, N.; Jensen, D. R.; Fujii, H.; Zhao, Y.; Lumba, S.; Santiago, J.; Rodrigues, A.; Chow, T. F.; Alfred, S. E.; Bonetta, D.; Finkelstein, R.; Provart, N. J.; Desveaux, D.; Rodriguez, P. L.; McCourt, P.; Zhu, J. K.; Schroeder, J. I.; Volkman, B. F.; Cutler, S. R., Abscisic acid inhibits type 2C protein phosphatases via the PYR/PYL family of START proteins. Science 2009, 324 (5930), 1068–71.

42. Lin, Z.; Li, Y.; Zhang, Z.; Liu, X.; Hsu, C. C.; Du, Y.; Sang, T.; Zhu, C.; Wang, Y.; Satheesh, V.; Pratibha, P.; Zhao, Y.; Song, C. P.; Tao, W. A.; Zhu, J. K.; Wang, P., A RAF-SnRK2 kinase cascade mediates early osmotic stress signaling in higher plants. Nat Commun 2020, 11 (1), 613.

43. Takahashi, Y.; Zhang, J.; Hsu, P. K.; Ceciliato, P. H. O.; Zhang, L.; Dubeaux, G.; Munemasa, S.; Ge, C.; Zhao, Y.; Hauser, F.; Schroeder, J. I., MAP3Kinase-dependent SnRK2-kinase activation is required for abscisic acid signal transduction and rapid osmotic stress response. Nat Commun 2020, 11 (1), 12.

44. Lin, Z.; Li, Y.; Wang, Y.; Liu, X.; Ma, L.; Zhang, Z.; Mu, C.; Zhang, Y.; Peng, L.; Xie, S.; Song, C. P.; Shi, H.; Zhu, J. K.; Wang, P., Initiation and amplification of SnRK2 activation in abscisic acid signaling. Nat Commun 2021, 12 (1), 2456.

45. Yoshida, T.; Fujita, Y.; Maruyama, K.; Mogami, J.; Todaka, D.; Shinozaki, K.; Yamaguchi-Shinozaki, K., Four Arabidopsis AREB/ABF transcription factors function predominantly in gene expression downstream of SnRK2 kinases in abscisic acid signalling in response to osmotic stress. Plant Cell Environ 2015, 38 (1), 35–49.

46. Peirats-Llobet, M.; Han, S. K.; Gonzalez-Guzman, M.; Jeong, C. W.; Rodriguez, L.; Belda-Palazon, B.; Wagner, D.; Rodriguez, P. L., A Direct Link between Abscisic Acid Sensing and the Chromatin-Remodeling ATPase BRAHMA via Core ABA Signaling Pathway Components. Mol Plant 2016, 9 (1), 136–147.

47. Wang, P.; Zhao, Y.; Li, Z.; Hsu, C. C.; Liu, X.; Fu, L.; Hou, Y. J.; Du, Y.; Xie, S.; Zhang, C.; Gao, J.; Cao, M.; Huang, X.; Zhu, Y.; Tang, K.; Wang, X.; Tao, W. A.; Xiong, Y.; Zhu, J. K., Reciprocal Regulation of the TOR Kinase and ABA Receptor Balances Plant Growth and Stress Response. Mol Cell 2018, 69 (1), 100–112 e6.

