## Supplementary material for "A Suspension Trapping–Based Sample Preparation Workflow for Sensitive Plant Phosphoproteomics": Figure S1, Figure S2, Figure S3, Figure S4, Table S1

**Table of content**

**Figure S1.** Comparison of the overlap of identified phosphopeptides from the SDS protocol and the Urea-SDS protocol.

**Figure S2.** Comparison of the ratio of quantifiable phosphopeptides from the Standard elution protocol and the Direct loading protocol.

**Figure S3.** Comparison of the ratio of missed cleavage from the S-Trap protocol and the Precipitation protocol.

**Figure S4.** The identified phosphopeptides of each replicate.

**Table S1.** The total number of identified phosphoproteins, phosphopeptides, and phosphorylation sites in the ABA-dependent phosphoproteomics experiment.

**Table S2.** Identified phosphopeptides and their intensities and localization probabilities in the ABA-dependent phosphoproteomics experiment (XLSX).

**Table S3.** The z-scored intensities of selected ABA signaling components (XLSX).

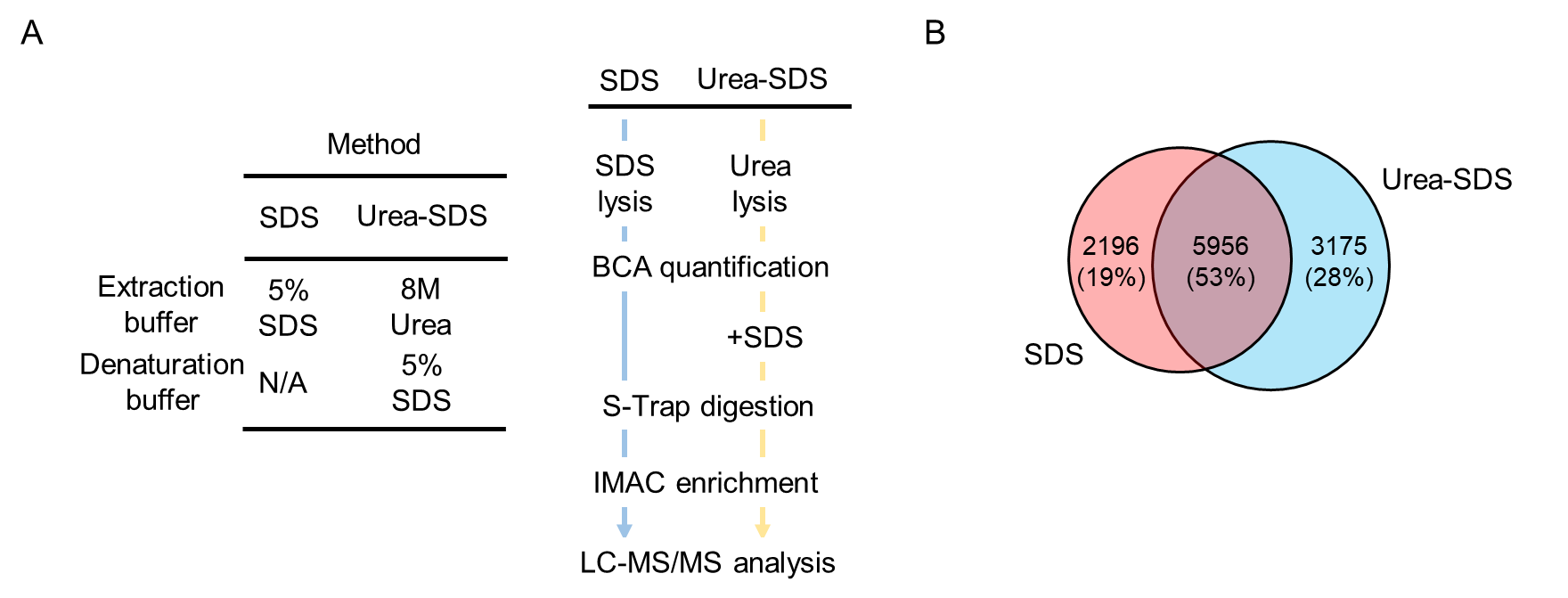

**Figure S1.** **Comparison of the overlap of identified phosphopeptides from the SDS protocol and the Urea-SDS protocol.** A. Left, the extraction buffer and protein denaturation buffer used in the SDS protocol and the Urea-SDS protocol. Right, schematic representation of two protocols. B. Venn diagram showing the overlap between identified phosphopeptides from the SDS protocol and the Urea-SDS protocol.

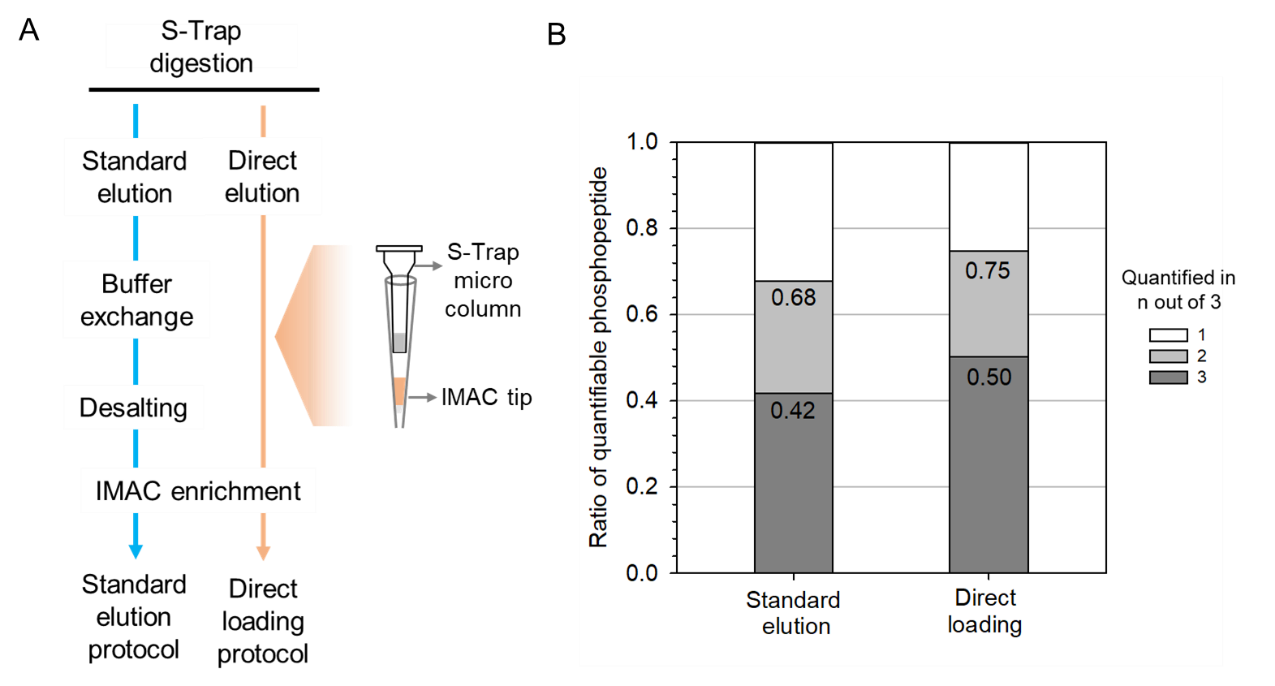

**Figure S2. Comparison of the ratio of quantifiable phosphopeptides from the Standard elution protocol and the Direct loading protocol.** A. Schematic representation of the protocols. B. The ratio of quantified phosphopeptides from triplicate replicates of each protocol.

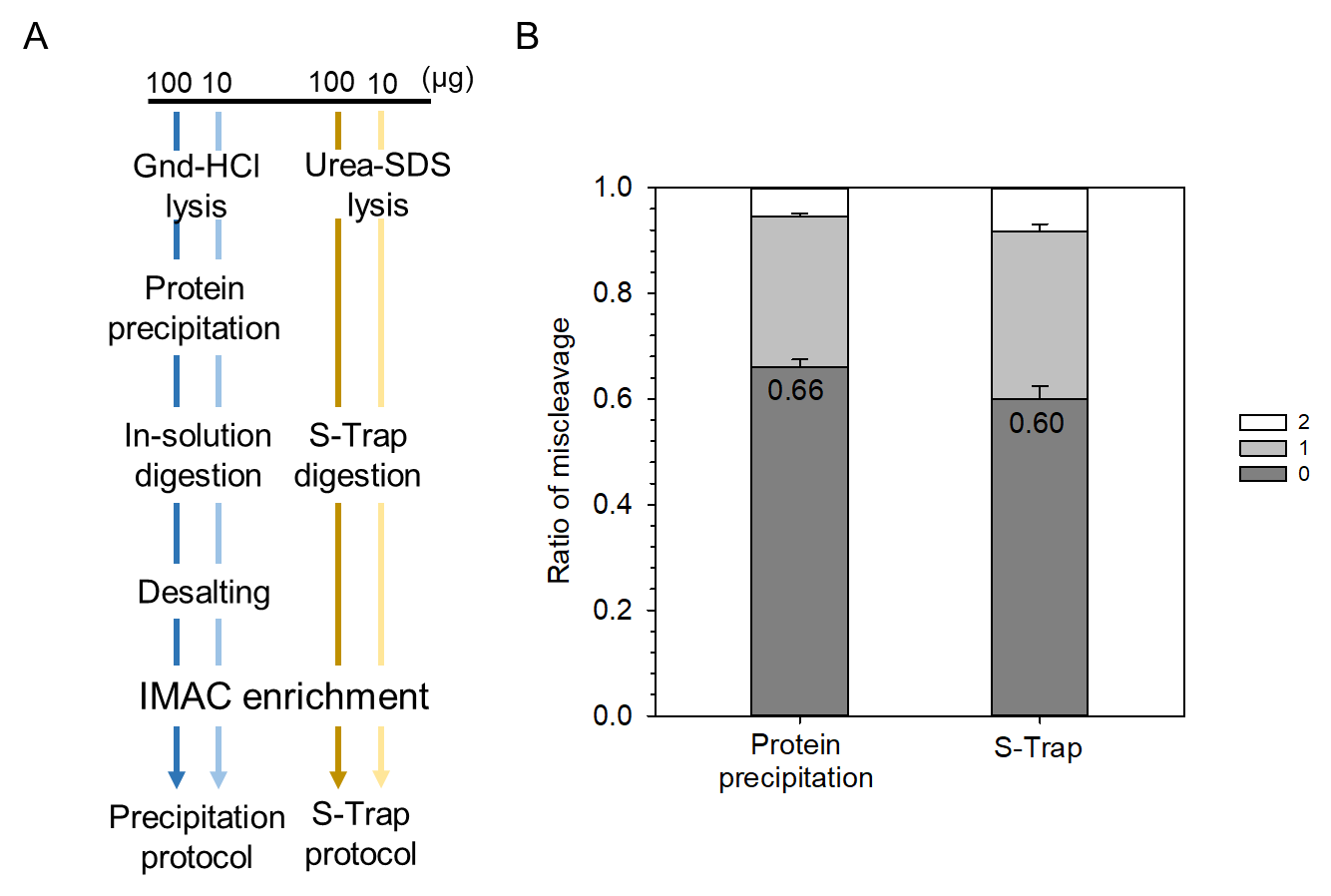

**Figure S3.** **Comparison of the ratio of missed cleavage from the S-Trap protocol and the Precipitation protocol.** A. Schematic representation of the protocols with low-input and high-input sample. B. Comparison of the ratio of tryptic missed cleavage between the protein precipitation protocol and the S-Trap protocol. Error bars, SD (n = 3).

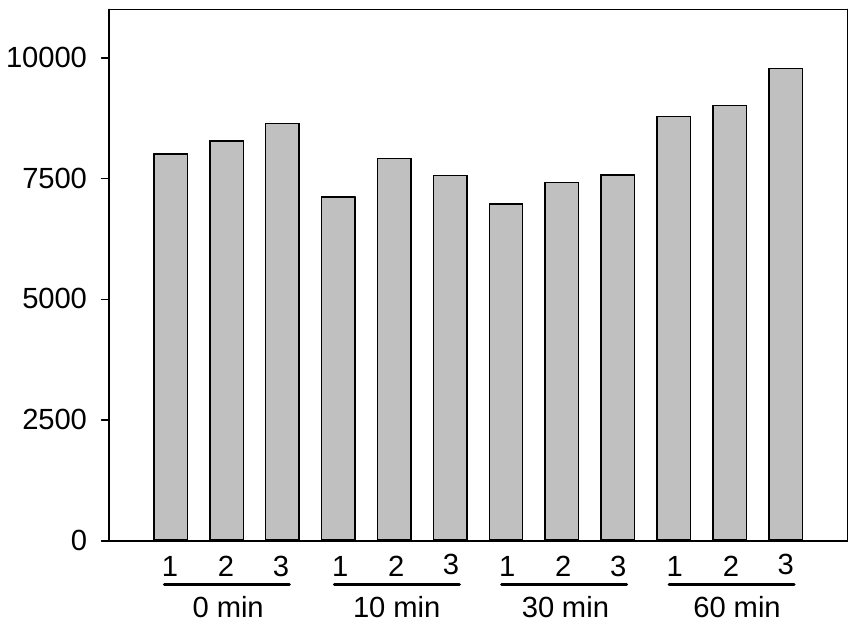

**Figure S4.** **The identified phosphopeptides of each replicate.**

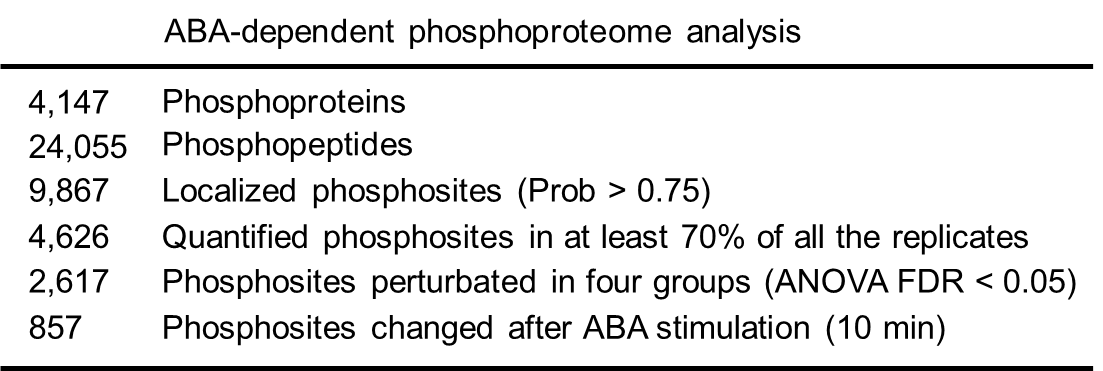

**Table S1. The total number of identified phosphoproteins, phosphopeptides, and phosphorylation sites in the ABA-dependent phosphoproteomics experiment.**
